# Peaceful behaviour: a strategy employed by an obligate nest invader to avoid conflict with its host species

**DOI:** 10.1101/587592

**Authors:** Helder Hugo, Paulo F. Cristaldo, Og DeSouza

## Abstract

In addition to its builders, termite nests are known to house a variety of secondary, opportunistic termite species, but little is known about the mechanisms governing the maintenance of such associations. In a single nest, host and intruder are likely to engage in intense conflict, due to their nestmate discrimination system. An intriguing question is how individuals cope with such a burden in the long term. Evasive behaviour has been previously suggested as a mechanism that reduces the frequency of encounters between non-nestmates. However, due to confinement imposed by the nests’ physical boundaries, it is likely that hosts and inquilines would eventually come across each other. Under these circumstances, it is plausible that inquilines would be required to behave accordingly to secure their housing. Here, we tested this hypothesis predicting that, once inevitably exposed to hosts, inquiline individuals would modulate their behaviour to circumvent conflict. While exploring the behavioural dynamics of the encounter between both cohabitants, we find evidence for an unusual lack of aggressiveness by inquilines towards hosts. Such a non-aggressive behaviour is characterised by evasive manoeuvres that include reversing direction, bypassing and a defensive mechanism using defecation to repel hosts. The behavioural adaptations we describe may play an intrinsic role in the stability of cohabitations between termite species: by reducing the costs of conflicts to both cohabitants, it may improve the chances for stable nest-sharing considerably.

## Introduction

Nature provides innumerable opportunities to observe animals coexisting (Tokeshi, 2009; Gravel et al. 2011), from migratory species temporarily interacting with local communities (Kays et al., 2015) to organisms establishing long-term, interspecific relationships (Wilson, 1988). Of particular interest, the latter group includes species that cohabit a single place and may, therefore, interact several times throughout their lifespan. These associations, or *symbioses* (*sensu* De Bary, 1878; see Oulhen et al., 2016), often represent excellent opportunities to investigate how organisms with independent life-histories end up sharing precisely the same place. For instance, while making decisions about permanent housing, some organisms opt for nests already built by a different species, avoiding costs with construction. In addition to providing shelter, nests may contain resources continuously renewed over time (e.g. food and water) and attract a variety of opportunistic organisms on the way. That seems to be the case of termite nests (termitaria), in which is possible to find an impressive richness and abundance of non-nestmates cohabiting with the original termite builders (Costa et al., 2009; Monteiro et al., 2017).

Although a wide variety of species have been found inside termitaria (Kistner, 1969, 1979, 1990; De Visser et al., 2008), in this paper we focus on a remarkably distinct case of nest cohabitation between builder and invader, that is, a host termite and a secondary, opportunistic termite species so-called *inquilines* (*sensu* Araujo, 1970). It is worthwhile mentioning, however, that inquilinism among termites should not be mistaken with that occurring in Hymenoptera. Commonly referred as *social parasites* (Nash & Boomsma, 2008), inquiline bees, wasps and ants tend to establish a close relationship with hosts and exploit their social behaviour intensively (Hölldobler & Wilson, 1990). In termites, though, inquilines are thought to be primarily associated with the nest’s physical structure itself, regardless of their association with host species (Shellman-Reeve, 1997; Marins et al., 2016). With proportionally smaller colonies and relatively low brood care (Korb et al., 2012) it is unlikely, although possible, that inquiline termites would deplete nest resources intensively, or exploit the host’s social structure, as reported in different inquiline ant species (Buschinger, 2009).

Framing precisely inquilinism among termites into the spectrum of symbiotic interactions (e.g. parasitism, commensalism) can be challenging. For instance, although a number of studies have provided relevant information on different host-inquiline systems (e.g. Collins 1980; Redford, 1984; Eggleton & Bignell 1997; Cunha et al. 2003; Costa et al. 2009; Darlington, 2011; Cristaldo et al. 2012, 2014; Florencio et al. 2013; Campbell et al., 2016; DeSouza et al. 2016; Rodrigues et al. 2018), it remains unclear which costs (if any) inquiline termite colonies impose to host species. Even so, it seems plausible that a community of termite species within a single nest would be an ideal scenario for the emergence of conflict. Because host termite species are known to respond aggressively towards a variety of nest intruders (Emerson, 1938; Shellman-Reeve, 1997), the confrontation would arise predominantly from encounters with non-nestmates. Aggressive behaviour seems to be, in fact, a default response of the soldier caste of termites towards non-nestmates (Noirot, 1970), with individuals engaging in endless fights while protecting their colonies (Binder 1987). Moreover, in addition to the typical agonism of soldiers, hidden aggression among termite workers has been reported for some species (Ishikawa & Miura 2012).

Curiously enough, as opposed to hosts, inquiline colonies may be found in the wild severely depleted in their contingent of soldiers (Cunha et al. 2003). The proportion of soldiers in some cases may account for less than one per cent of the colony (HH, pers. obs.). Relying on nest invasions to persist may, hence, represent a considerable risk for inquiline colonies, and an intriguing question is how cohabitation in such terms is even possible. Not surprisingly, previous works have tackled such an issue, suggesting proximate mechanisms that would allow inquilines and hosts to meet less frequently within the nest. In this regard, immediately after successful invasions, inquilines would establish themselves in the nest by decreasing chances of being noticed by hosts in the first place. Inquiline termites could achieve such an effect through various behaviours, including: (i) avoiding walking in galleries crowded by hosts (Grassé, 1986; Mathews, 1977); (ii) not conflicting with dietary requirements of the host (Miura & Matsumoto, 1997; Florencio et al., 2013); (iii) intercepting hosts’ chemical signals and using the information acquired to preclude encounters with hosts (Cristaldo et al., 2014, 2016a); and (iv) keeping the colony isolated from hosts by changing the nest structure (e.g. building their own galleries and sealing chambers: HH, pers. obs.). Although functioning through independent mechanisms, these behavioural strategies seem to coincide in a single outcome: by preventing direct contact, inquilines reduce the frequency of encounter with hosts.

As efficient as it may seem, however, while such strategies could potentially attenuate conflictual events, they would not entirely prevent encounters from happening. For most inquiline species (including the one studied here), there is no evidence yet of colonies exiting nests after they break in, neither for nest defence nor for foraging. The only known exception is the winged reproductive caste that leaves the nest during swarming (Matsuura, 2010). These facts together suggest that there is an associated probability of interspecific encounter to be considered. Besides, the confinement imposed by the nest’s physical boundaries would keep individuals locally restricted and bound to meet in the long-term. Under these circumstances inquilines would be required to behave accordingly, for instance, mitigating detrimental consequences of aggressive encounters with nest owners.

Although hinted in the past, this intuitive, theoretical prediction was never directly tested, and little is known about host-inquiline dynamics within the nest, or to what extent inquiline strategies are sufficient to cope with the menace of imminent confrontation with hosts. This information could provide important clues about how these cohabitations hold in nature. A conservative approach to this issue would sustain that inquilines should replicate, at the individual level, the evasive behaviour they exhibit as a group. In this context, one would expect the strategies to avoid conflict (highlighted above) to be mere consequences of a non-threatening posture exhibited by inquiline individuals. Here, we tested this hypothesis predicting that, once inevitably exposed to hosts, inquiline individuals would adopt a non-aggressive posture and modulate their behaviour to a less threatening profile. As a result, the colony would be able to reduce conflict with nest owners collectively. Such an assumption would imply that inquiline individuals should weaken conflict escalation by (i) being lethargic and minimising encounters with hosts and (ii) exhibiting low aggressiveness by avoiding either initiating or retaliating attacks.

To test these assumptions, we observed in detail the behaviour of an obligate inquiline termite, *Inquilinitermes microcerus* Silvestri (1901) (Termitidae: Termitinae), in the presence of its host termite, *Constrictotermes cyphergaster* Silvestri 1901 (Termitidae: Nasutitermitidae). We exposed species to each other under two different experimental scenarios: (i) in closed arenas, as to keep individuals locally restricted and favour host-inquiline encounters; and (ii) in open arenas, where inquilines had a chance to flee from hosts. By compiling full ethograms for the encounter between *I. microcerus* and *C. cyphergaster*, we add new information to the current knowledge on nest-sharing termite species. These descriptions highlight relevant aspects to consider while studying the underlying mechanisms of coexistence between species living in environments circumscribed by discrete physical barriers. Furthermore, we argue that the behavioural profiles here described lend support to the notion of inquilines as peaceful guests, contributing to the growing view of conflict-avoidance as an effective strategy to coexist in harsh environments.

## Methods

### Biological model

The termite *C. cyphergaster* (hereafter, host) is a Neotropical species widely distributed in South America (Mathews, 1977; Krishna et al., 2013) known to forage at night in exposed columns and without the protection of covered galleries (Moura et al., 2006). In this species, nest foundation starts on the ground with a royal couple, and after reaching a certain size, colonies migrate to the trees, where they establish typical arboreal nests (Vasconcellos et al., 2007). At this phase, it is usual to find colonies of *I. microcerus*, a secondary opportunistic termite species that inhabit the nests (hereafter, inquilines). Such a suggestive name as *Inquilinitermes* has its reasons: inquilines seem to be unable to build nests by their own (Emerson, 1938; Mathews, 1977), being found so far exclusively within host nests. Although it remains unclear how exactly nest invasion occurs, there seems to be a critical nest volume above which inquiline colonies are more likely to be found within host nests (13.6 L, see Cristaldo et al., 2012). The nest’s size seems to indirectly affect inquilinism in termites, being negatively related to defence rates (DeSouza et al., 2016). Besides, while evaluating populational parameters of nests containing inquiline colonies, Rodrigues et al. (2018) reported a negative correlation between the number of individuals and the proportion of soldier/workers. Compared to hosts, inquiline colonies are much smaller in size, but still easily detectable due to a characteristic dark lining covering their galleries (Cunha et al., 2003; Cristaldo et al., 2012; Florencio et al., 2013). Inside the nest, inquiline colonies are often associated with chambers filled with a black material, hypothesised in the past as waste dumped by hosts (Emerson, 1938), but still of unknown origin.

### Study site and collection

To carry out experiments, 27 host nests with inquiline colonies were collected from two locations in the Brazilian Cerrado (Ratter, Ribeiro, & Bridgewater, 1997): 15 nests collected near the municipality of Sete Lagoas (19º27’57”S, 44º14’48”W) in July 2012; and 12 nests collected near the municipality of Divinópolis (20º08’20”S, 44º53’02”W) in January 2015. Both sites, located in Southeastern Brazil (State of Minas Gerais), have climate resembling savannas (Aw, in Köppen-Geiger classification), and are subjected to an equatorial climate with dry winters (Aw) (Kottek et al., 2006).

### Experimental design

In order to access behavioural profiles at host-inquiline encounters, cohabitants were taken from their nests, acclimatised for 30 minutes in separate containers, and then gathered in arenas for video recording (Fig. 1). Experimental arenas consisted of plastic Petri dishes (Ø 53mm) lined with paper (Whatman N° 1), and video-samples were taken with a digital camera (Nikon D300S, 720p, 25fps). All videos were recorded under visible light, and the room temperature was controlled between 23°C and 25°C.

**Figure 1.**
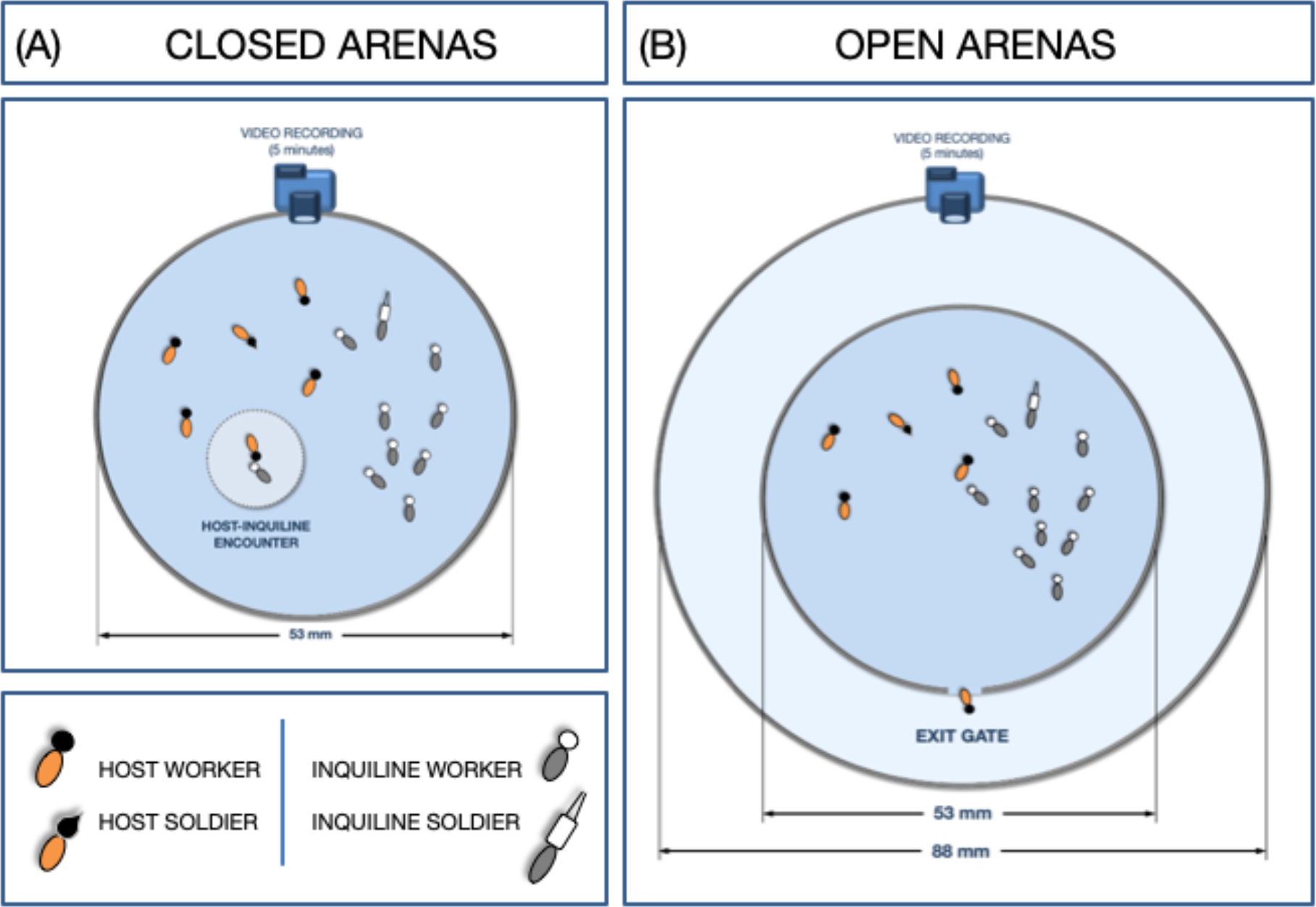
Arena settings for video recording: (A) Closed arenas, (B) Open arenas. We recorded video-samples of five minutes (300 seconds) for focal animal observation, which was carried out by observers using a 14” LED-LCD 1080p screen; an example of a heterospecific encounter is highlighted by a lighter circle within the closed arena (A). The internal area had a diameter of 53mm. The external area had a diameter of 88mm. A gate consisted of a single opening with diameter of 3.5mm on the arena wall connecting internal and external areas.

We designed two experiments to test our predictions. In the first experiment (Fig. 1A), host and inquiline individuals were mutually confined in closed arenas, a locally restricted condition intentionally designed to improve the chances of an encounter between the species. In the second experiment, hosts and inquilines were gathered in arenas mostly identical to those used in the first setup, except for the presence of an exit gate. This gate consisted basically of a single opening (Ø 3.5mm) on the arena wall, giving access to an external circular area (Ø 88mm) encompassing the inner one (Ø 53mm). This second experiment was conducted to inspect whether inquilines would (i) remain idle or (ii) move away when given a chance to flee from host aggressions (Fig. 1B). The latter response could potentially lead to spatial segregation between species, a result that would be in line in field observations.

We defined two treatments using open arenas to test whether the presence of inquilines would affect how hosts explore the space available: (i) open arenas containing host and inquilines; and (ii) open arenas containing only hosts, as a control. Experiments were conducted with individuals kept under optimal density (0.12; for details, see Miramontes & DeSouza, 2008) and in a worker-to-soldier ratio similar to that found in natural conditions (Cunha et al., 2003). Experimental groups, therefore, contained: (i) one soldier and four workers for hosts and (ii) one soldier and nine workers, for inquilines. Individuals composing a given experimental group were never present in a second trial, as to avoid interference from prior contact with non-nestmates.

### Behaviour annotation and observational protocol

To determine observable behaviours of relevance to our scope, we performed the following procedure: before the main experiments, preliminary observations were taken as to detect behaviours possibly performed by individuals. Video-samples used to perform these observations were never reused in the main experiments. At this phase, we spent efforts to describe as many behaviours as possible. Because extensive behavioural descriptions may contribute to misleading observation (Lehner, 1998), we created a flowchart with simple, straightforward labels hierarchically organised (Fig. 2). Observers used this diagram and a list of short behavioural descriptions (Table 1) as a reference for annotation.

**Table 1.**
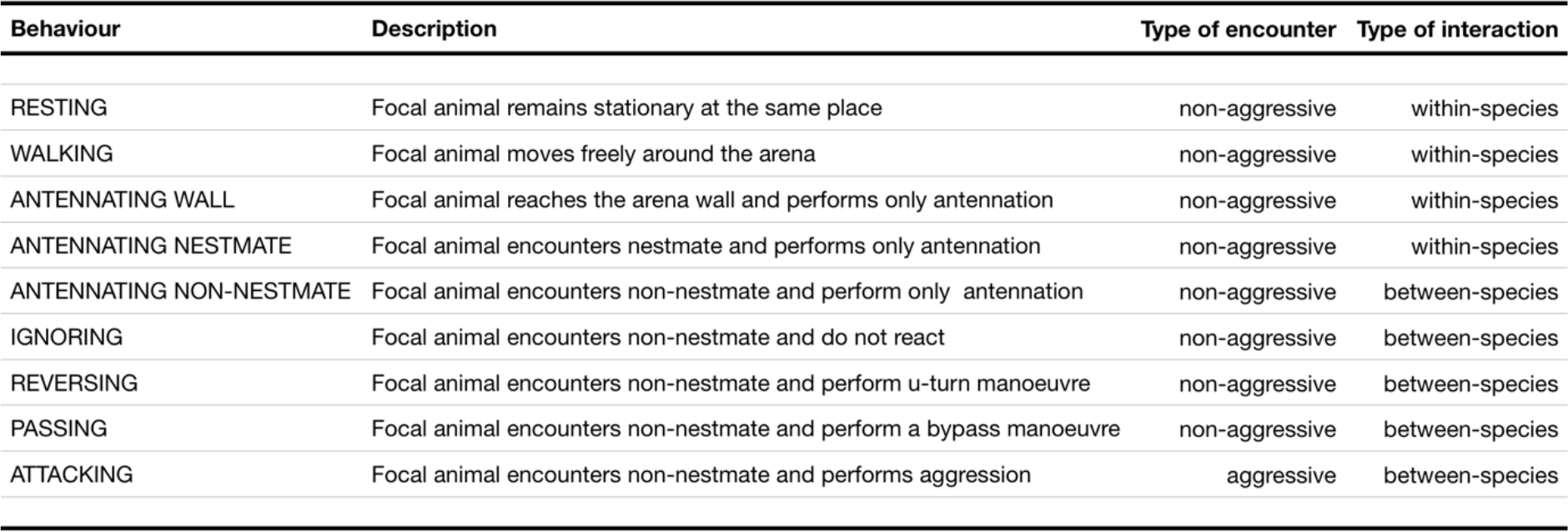
Behavioural description based on preliminary observations. We defined nine observable behaviours of relevance to our scope using ten additional video-samples. For statistical analysis, we classified each behaviour in two ways: (i) either as within-species or between-species; and (ii) either as aggressive or non-aggressive.

**Figure 2.**
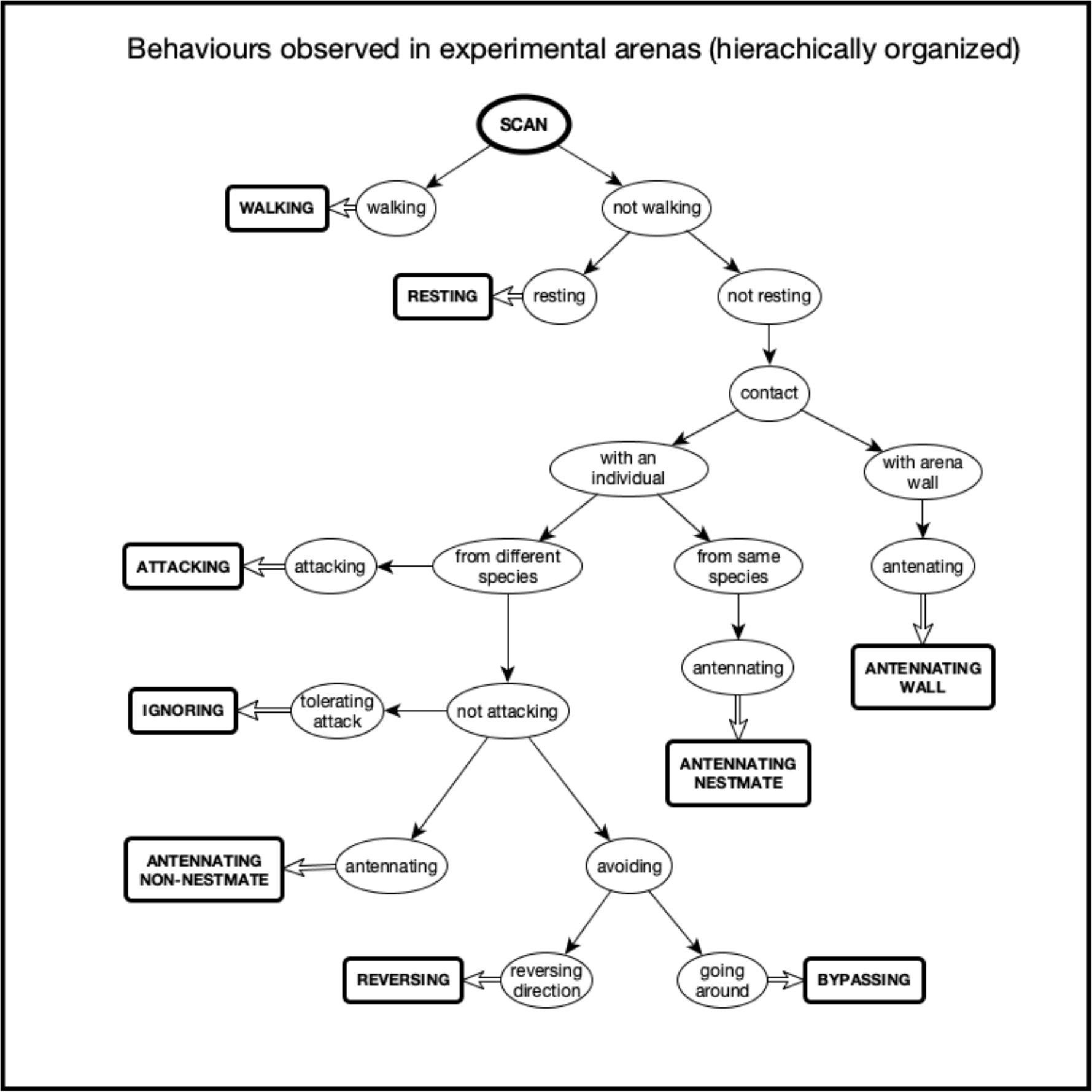
Behavioural flowchart used for annotation. Labels are hierarchically organised to allow stepwise classification of behaviours. We defined nine observable behaviours in preliminary observations. Combined to a behavioural description presented in Table 1, this flowchart served as a reference for annotation.

We adopted focal animal sampling (Altmann, 1974) with observations taken from video-samples to capture interactions between hosts and inquiline individuals. We recorded a total of 20 video-samples of five minutes (10 using closed arenas, 10 using open arenas). Behaviours performed by host workers (HW), host soldiers (HS), inquiline workers (IW) and inquiline soldiers (IS) were annotated for each video-sample using the flowchart described above. For each one of these categories, an individual was arbitrarily selected for focal observation (hereafter, focal animal). Using a 14” LED 1080p screen to watch video samples, we took three-seconds observations (hereafter, scans) for each focal animal. Scans were taken at regular time intervals of 10 seconds, indicated to observers by scheduled sound signals. This method provided 31 scans *per* focal animal for each video-sample. Finally, we organised all behavioural annotation in files including all relevant information (e.g. observer, date and time of recording, room temperature)data analyses.

### Measuring aggressiveness and host-inquiline interactivity

To measure host’s and inquiline’s aggressiveness in closed arenas, we classified behavioural observations into two mutually exclusive categories regarding the type of encounter: (i) aggressive encounter, when focal animals encountered non-nestmates and performed aggression (i.e. attack); and (ii) non-aggressive encounter, when focal animals interacted with non-nestmates but did not perform aggression (i.e. antennating non-nestmate, reversing, bypassing, ignoring). To measure interactivity between hosts and inquilines in closed arenas, we classified behavioural observations into two mutually exclusive categories regarding the type of interaction: (i) within-species, when focal animals performed actions either by themselves (i.e. resting, walking, antennating wall) or with nestmates (i.e. antennating nestmate); and (ii) between-species, when focal animals performed actions after establishing physical contact with non-nestmates (i.e. antennating non-nestmate, ignoring, bypassing, reversing, attacking).

### Assessing behavioural profiles

To analyse the influence of specific behaviours in the general profile of hosts and inquilines, we developed a network analysis using the free software yEd Graph Editor version 3.14.4 (yWorks, 2015). To build graphs for each caste, we performed the following procedure: using behavioural sequences extracted from annotations, we constructed adjacent matrices containing the behavioural change for each caste (Supplementary Material, **Table S2**), which later was imported to yEd to draw the graphs. As typically done in standard network analysis (Newman, 2003), graphs consisted of networks of nodes linked by connecting edges (*i.e*. directional arrows). In our case, however, nodes represented specific observable behaviours executed by individuals, whereas connecting edges represented behavioural changes from a given behaviour to another one. That is, if individuals changed from rest to walking behaviour, the behavioural change annotated would be rest-walk. With nine observable behaviours defined in our scope (Table 1), 81 types of behavioural change could be possibly observed. With the constructed graphs, we calculated centrality measures (Freeman, 1978) using the number of incoming connecting edges for each node (Brandes & Erlebach, 2005). Then, using the calculated centrality scores, we adjusted the size of nodes to visually represent the degree of influence exerted by each behaviour upon profiles (i.e. the larger the size of a node, the higher its influence on the network).

### Ethogram validation

We adopted a procedure suggested by Dias et al. (2009) to validate our ethograms. Following this approach, we used behavioural accumulation curves (BAC) to assess an optimal balance between (i) effort with sampling and (ii) ethogram completeness. A minimum of 250 independent observations would be required to efficiently capture a total of ten observable behaviours (Fig. 3). In our study, we extrapolated this number and performed 1240 discrete observations for the nine observable behaviours previously defined (that is, 31 scans × 2 castes × 2 species × 10 replicates = 1240 scans).

**Figure 3.**
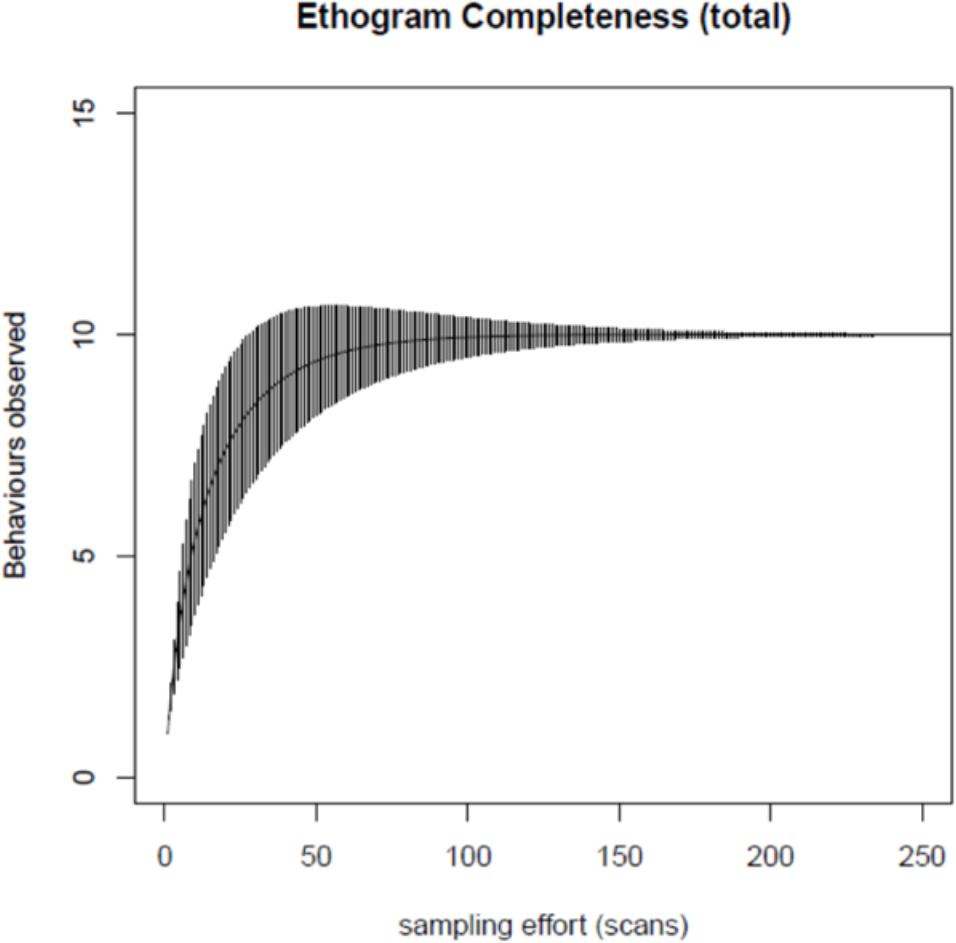
Ethogram completeness using Behavioural Accumulation Curves (Dias et al., 2009): the X-axis represents sampling effort, that is, the number of scans performed to observe all behaviours (in our case, 31 scans × 2 castes × 2 species × 10 replicates = 1240 scans); the Y-axis represents the accumulative number of behaviours experimentally observed in trials.

### Statistical analyses

We performed the statistical analyses in R, version 3.5.2 (R Development Core Team, 2018) using Generalized Linear Modelling (GLM) under Binomial errors with log-link. As a conservative approach, the significance of treatments was accessed using the following procedure: we compared complex models to simpler ones achieved by combining treatment levels (Crawley, 2012). When simplification did not provoke significant changes, simpler models were accepted, and the combined treatments were considered equivalent to each other. We then submitted adjusted models to a residual analysis as to check the suitability of the modelling equation and normality of error distribution. If required, error distribution was adjusted using Quasi-binomial distribution. In all tests conducted, we considered an α = 0.05 to assess statistical significance.

## Results

### Inquilines suffered attacks from hosts but responded with low aggressiveness

In closed arenas, the proportion of aggressive interactions initiated by hosts when encountering inquilines was significantly higher than the proportion of non-aggressive interactions (*GLM*; F_1,98_=16.72, *P<*0.001; Fig. 4). As expected, caste was determinant in the type of aggression inflicted by individuals. While host workers physically injured inquilines by biting them in several portions of their softy bodies, host soldiers frequently adopted an agonistic display, characterised by an abrupt movement with stretched antennae. In *C. cyphergaster*, soldiers present a snout-like protuberance in their head that contains a frontal gland. Such an apparatus produces a mixture of terpenoids, often used against targets in defensive actions (for details, see Cristaldo et al., 2015). In this study, however, we were not always able to detect whether an agonistic display was followed by chemical spray.

**Figure 4.**
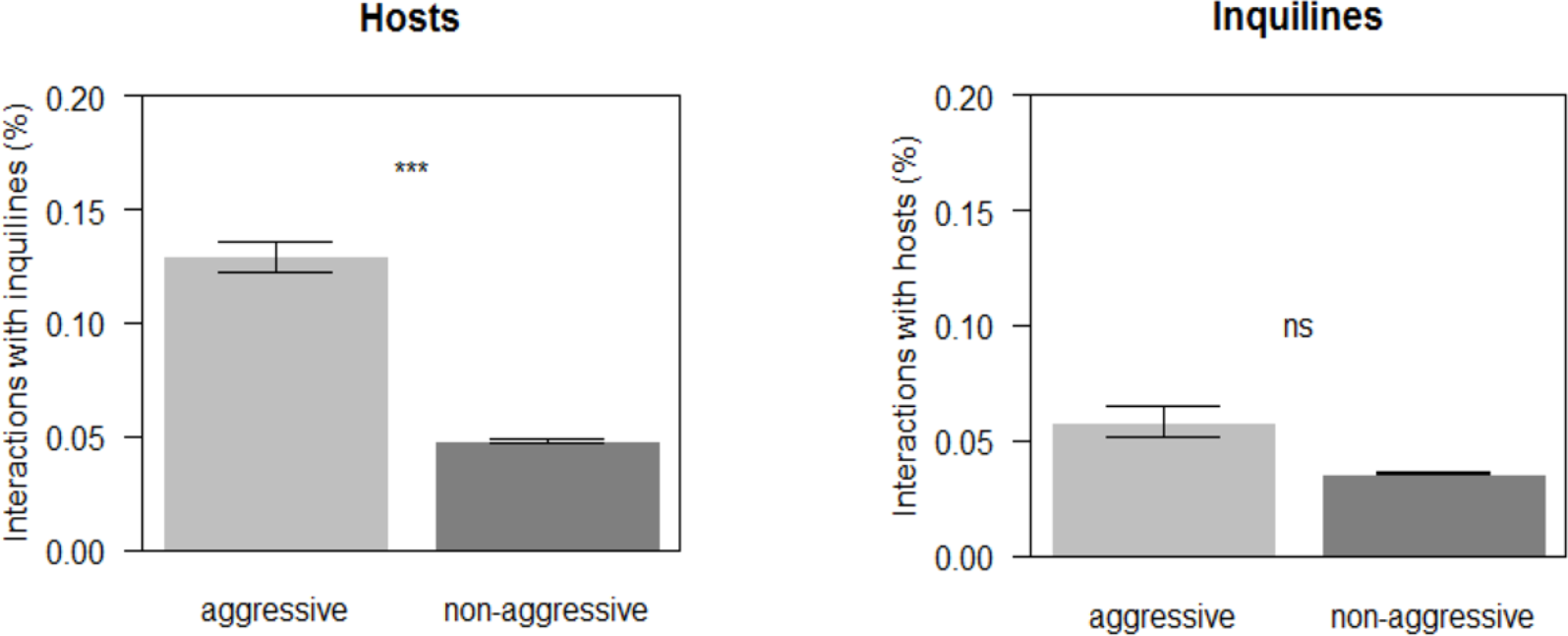
Interactions observed in closed arenas for hosts, *C. cyphergaster* (right), and inquilines, *I. microcerus* (left). Proportions were calculated by the number of aggressive and non-aggressive interactions observed, divided by the total number of observations taken from video-samples (N=10). Behaviours that do not preclude interaction (i.e. resting, walking and antennating wall) are not represented. For this reason, frequencies do not sum up to 100%. Light bars: aggressive interactions; Dark bars: non-aggressive interactions.

When attacked by hosts, the proportion of aggressive reactions initiated by inquilines was not significantly higher than the proportion of non-aggressive interactions (GLM; F_1,98_=1.74, P=0.18; Fig. 4). When threatened, or even severely injured by hosts, inquiline workers never retaliated (attacking; Fig. 5). Instead, individuals were more likely to adopt evasive manoeuvres and quickly divert from aggressors. These actions occurred immediately after an active contact with host individuals was established, and included behaviours that avoided the opponents (reversing, bypassing; Fig. 5). Besides escaping from host threats, inquiline workers also performed ignoring behaviour. In this case, individuals actively touched by hosts did not react to such a stimulus, remaining completely stationary (ignoring; Fig. 5).

**Figure 5.**
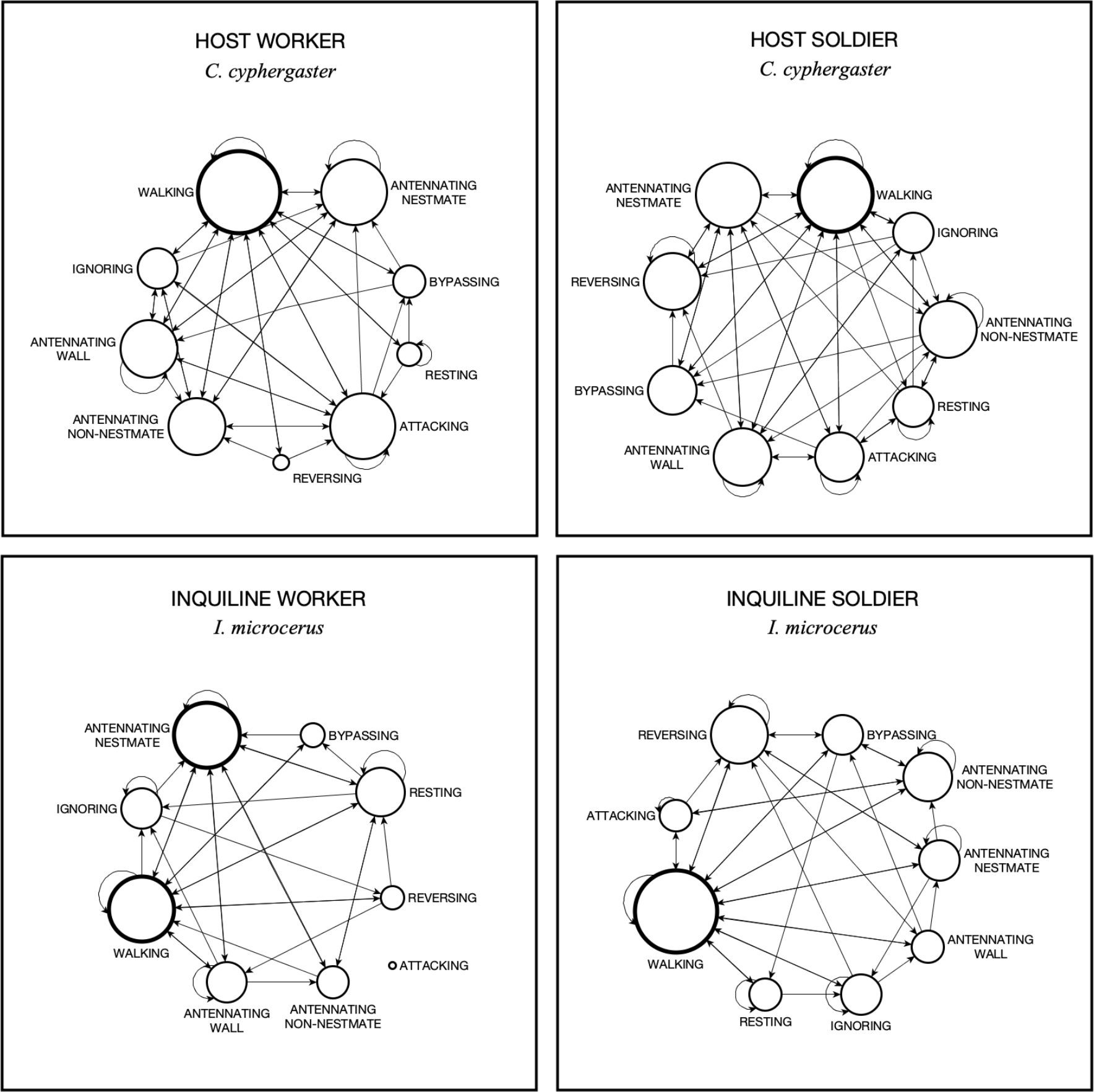
Behavioural profiles observed for each caste. Nodes represent behaviours performed by individuals, whereas connecting edges (arrows) represent behavioural changes occurred from one behaviour to another. Behaviours with the highest influence on the network are highlighted with thicker node contours. Node size was adjusted using calculated centrality measures to visually represent the degree of influence exerted by each behaviour upon the profiles. Obs.: For inquiline worker, attacking never happened and, therefore, this behaviour is not connected to the network. A version containing calculated scores is included in the Supplementary material (Figure S1).

The behaviour of inquiline soldiers was an exception to such a lack of aggressiveness among inquilines. As opposed to inquiline workers, inquiline soldiers performed aggression in retaliation to host assaults (Supplementary Material, **Video S1**), even though this occurred not so frequently (attacking, Fig. 5). Aggression performed by inquiline soldiers consisted of snapping attacks, a sudden release of slender mandibles pressed against each other producing powerful strikes over opponents (Supplementary Material, **Video S2**).

### Inquilines interacted little with hosts even when locally restricted

Inquilines exhibited low interactivity with host individuals, even when locally restricted and presumably more prone to meet, such as in closed arenas. The proportion of between-species observations was considerably lower than the proportion of within-species observations (*GLM;* F_1,178_= 71.73, *P<* 0.001). Besides, regarding the behavioural change, inquiline workers exhibited a loop between resting, walking and antennating nestmate, all behaviours with no contact with the host (**Figure S3**). The **Table S1** of Supplementary Material contains absolute numbers for between- and within-species observations for each one of the castes.

### Hosts were active in arenas, while inquilines were lethargic

When placed in closed arenas, host individuals performed antennation on the arena wall more frequently than inquiline individuals (GLM; F_1,78_= 4.73, *P<* 0.005), an indication that hosts could be possibly attempting to broaden their patrolled area. We confirmed this suspicion with results from the second experiment, with open arenas: host individuals quickly moved to the external area passing through the gate as soon as they found it. Either in the presence or absence of inquiline individuals, there was no difference in the mean time spent by host individuals to leave the internal area (19.93±3.56 seconds, F_1,9_=0.34, *P=* 0.57), meaning that inquilines did not threaten hosts. Inquilines, in turn, were more prone to remain stationary and never left the internal area, a result that seem to confirm the putative lethargic behaviour of inquilines.

### Inquiline’s defecation prevented host aggression

We observed an unexpected response among inquiline workers: when threatened by hosts, inquiline workers deposited faecal pellets always towards the direction from which they suffered threats (Supplementary Material, **Video S3**). Rather than usual defecation, this behaviour seemed to be elicited by host aggressions in a particular way: when receiving attacks from backwards, individuals immediately placed faecal pellets in front of aggressors and escaped forward. Threats coming from any other direction, however, triggered a slightly different response: before defecating, individuals first adjusted their posture accordingly, placing themselves in a way that they could quickly drop faecal pellets in front of the aggressors. Only after such a move, inquiline workers defecated and escaped forward. We observed this behaviour 33 times, and in all occurrences, faecal pellets immediately prevented inquilines of being chased of receiving further attacks from aggressors. Although we did not measure whether inquiline faeces have a repellent effect over hosts, it was evident in our recordings that areas containing faeces were less visited by host individuals (Supplementary Material, **Video S3**).

### Caste types showed unique behavioural profiles

We found striking differences when comparing the behavioural profile of hosts and inquilines (Fig. 5). Visual representations obtained from network analyses revealed unique configurations for each of the castes analysed. For all caste types, walking was the behaviour with the highest centrality score (walking; Fig. 5), that is, with the highest influence. The only caste type that presented two behaviours equally influential to the network was that of inquiline workers. In this group, besides walking, conspecific antennation also reached the highest centrality score (antennating nestmate; Fig. 5). As highlighted above, because inquiline workers never performed an attack, this behaviour presented the lowest centrally score and did not connect to the network (attacking; Fig. 5).

## Discussion

In environments where individuals are constantly surrounded by potential aggressors, the evolution of a “peaceful behaviour” may appear, at first, counter-intuitive. Our results demonstrate, however, that for inquilines (*I. microcerus*) non-aggressive behaviour is a valid strategy that mitigates detrimental consequences of unexpected encounters with hosts (*C. cyphergaster*). More important, it seems to secure housing for inquiline colonies within host nests in the long-term. Evasive behaviour by inquilines has been previously suggested as one of the proximate causes in inquilinism, as it reduces the frequency of encounter between colonies. Here, besides providing substantial behavioural data supporting this idea, we showed that once inevitably exposed to hosts, inquiline individuals can modulate their behaviour to a less threatening profile and circumvent confrontation. As compared to other termite species, this suggests a degree of adaptation towards a more flexible behaviour which could, in turn, strongly favour cohabitation.

### The behavioural adaptations of a peaceful guest

A set of behaviours seem to support our interpretation of inquilines as peaceful guests within host nests. First, when encountering host individuals, inquilines suffered several attacks but did not react with the same level of aggressiveness. Lack of aggression was markedly evident among inquiline workers. This caste not only never performed a single attack during our experiments but also managed to move away from aggressors with evasive manoeuvres (reversing, bypassing; Fig. 5). The same was not observed for inquiline soldiers, who did retaliate host attacks with snapping (attacking; Fig. 5). It is worth mentioning, however, that soldiers are rare in natural colonies of *I. microcerus* (Cunha et al., 2003), and sometimes even absent (HH, pers. obs.). Thus, it is unlikely that the aggressive behaviour of a minority would contribute to increase the species aggressiveness substantially. Although we did not find evidence pointing in such a direction, we acknowledge the necessity of more studies evaluating the interplay between worker’s and soldier’s behaviour in field conditions, something that was clearly beyond the scope of the present work.

A second behaviour linked to the levels of aggressiveness reported (Fig. 3) was the reduced mobility of inquilines. Consequently, the interaction between hosts and inquilines in arenas was limited. In open arenas, only hosts moved to the external area over time, whereas inquilines remained quiet in the inner portion. Such spatiotemporal segregation could be a direct consequence of the behavioural profile of hosts. As we have shown, walking was a commonly performed behaviour among all castes, but at the same hosts spent less time remaining stationary in the same place (resting, Fig. 5). Theoretically, as they walk more intensively and explore sites more efficiently, gates would be more readily found. In this regard, the presence or absence of non-nestmates in arenas did not affect the time spent by hosts to pass through the gate and access the external area. This result indicates that inquilines did not necessarily triggered the collective motion of hosts to the external area. Dynamics of both collective behaviour and environment have been suggested to regulate group-level properties in ants (Gordon, 2019). In termites, some studies have explored principles of collective behaviour (Sumpter, 2006) using agent-based models to understand self-organisation of groups, from nest construction processes (Deneubourg, 1977) to aspects of social facilitation (Miramontes & DeSouza, 1996; DeSouza et al., 2001). Still, for nest-sharing termite species, to what extent individual behaviour shape collective motion patterns, remains a topic to be fully understood.

A third component that seemingly affected the amount of aggression we observed in arenas was defecation by inquilines. Presence of faecal pellets shortened host-inquiline contact in virtually all occasions, and consequently, host attacks towards inquilines were less frequent. This result indicates that faeces may improve evasion by preventing host aggressions. In fact, defecation as an evasive mechanism is not exclusive of *I. microcerus*, being first described for termites by Coaton (1971) in *Skatitermes* sp. (Termitidae: Apicotermitinae). Such a defensive behaviour may have important implications for cohabitation: if faeces indeed repel hosts, single pellets placed in narrowed galleries throughout the nest could prevent host contact in a very efficient, inexpensive way. Besides, it is possible that while placing the pellets, *I. microcerus* would be spreading their scent throughout the entire nest, making it harder for hosts to locate the core of their colonies. Accordingly, while studying the cohabitation of another host-inquiline pair (*C. cavifrons* and *I. inquilinus*, respectively), Jirošová et al. (2016) showed that walls from the inquiline portion of nests contain levels of C12 alcohols, a repellent for host individuals. According to these authors, chemically mediated spatial separation of hosts and inquilines may aid to avoid conflict. Among other non-related groups, such as the cuckoo bumblebee, repellent odours are known to reduce host attacks (Lhomme et al., 2012), suggesting that this is an effective mechanism across taxa.

### The meaning of an interspecific encounter

The non-aggressive behaviour observed among inquilines raises the question of whether such a strategy would be useful within the nest. After all: are encounters with the host species a real threat for inquiline colonies? We provide evidence that there are, indeed, detrimental consequences of encountering hosts. As mentioned before, aggression among hosts was performed not only by soldiers but also by workers. (attacking, Fig. 5), which seems to indicate that defence would be integrated between castes. While host soldiers attack individuals spraying chemicals and provoking disruptive reactions, host workers provided with functional mandibles, are the ones who inflict the physical damage. In *C. cyphergaster*, terpenoids sprayed from the frontal gland of soldiers function as an effective alarm pheromone (Cristaldo et al. 2015). Thus, once a target is sprayed, it recruits nestmates to converge upon the site and deploy themselves around it (Eisner et al., 1976). In such a harsh environment, where virtually all individuals are potential aggressors, it is plausible that a peaceful behaviour, rather than a costly aggressive profile, could be a simpler alternative solution. All in all, as compared to a belligerent set of behaviours, a non-threatening profile would demand less elaborated actions, plausibly resulting in lower activity and reduced probability of interspecific encounter.

### The mechanism of conflict avoidance

Behaviours preventing confront escalation are widespread. When attacked by host ant workers, for instance, parasite ant queens do not react aggressively and, instead, quickly move towards the fungus garden remaining quietly there (Nehring et al., 2015). In another typical social parasite, *Maculinea rabeli* (Lycaenidae), larvae individuals are known to suppress aggression from their host ants by mimicking aspects of the brood’s pheromone (Akino et al., 1999; Pierce et al., 2002). Among bees, changes in the host’s behaviour towards non-aggressive types have been also reported: in the presence of the cuckoo bumblebee *Bombus vestalis*, host colonies decrease worker aggressiveness towards alien individuals, possibly due to changes in the host worker’s discrimination (Lhomme et al., 2012). Examples of aggressiveness being affected by external factors are not exclusive to social insects, extending to other invertebrate and vertebrate groups (Aureli et al., 2002; Baan et al., 2014; Gobush & Wasser, 2009; Thierry et al., 2008). In termites, aggressiveness may depend on factors such as diet (Florane et al., 2004), caste ratios (Roisin et al., 1990), nestmate recognition (Delphia et al., 2003; Haverty & Thorne, 1989), group composition (Haverty & Thorne, 1989), territoriality (Adams & Levings, 1987; Levings & Adams, 1984) and resource availability (Cristaldo et al., 2016b). Even inter-colony aggression, presumably more predictable due to higher relatedness, is not always consistent (Binder, 1987). Species may exhibit behavioural plasticity (Ishikawa & Miura 2012), responding aggressively in some cases (Su & Haverty, 1991), and lacking aggression in others (Delaplane, 1991; Neoh et al. 2012). Altogether, these reports indicate that it is possible to have scenarios in which termite species adopt low aggressiveness profile, rather than the typical aggressiveness observed among the group.

The symbiosis between *C. cyphergaster* and *I. microcerus* is a case of obligate inquilinism, meaning that at least for inquilines, nest-sharing has become mandatory (Shellman-Reeve, 1997). Evolutionary costs and drawbacks of such a specialisation by inquilines remain to be assessed, although the benefits associated with nest invasion seem to be straightforward: nest invaders are not required to spend time and energy building their own home. At the same time, being nest construction a demanding, costly process (Korb & Linsenmair, 1999), one would expect such inquiline invasions to be not strictly in the interest of hosts. In this sense, it would be reasonable to think of a scenario in which hosts would endeavour to detect inquilines, whereas inquilines would try to go unnoticed by hosts. Under such driving forces, it is possible that an evolutionary arms race would take place (Dawkins & Krebs, 1979), leading hosts and inquilines to reach well-adjusted behavioural profiles. In doing so, both cohabitant would become highly specialised in their neighbour (Kilner & Langmore, 2011).

### Cohabitation and conflict

Cohabitation goes way beyond the “living in overlapping spaces”. Instead, it is a result of multiple interactions over time. Whether interactions contribute for the emergence of stable relationships depends on the consequences mutually inflicted by the parties involved. With our approach, we presented findings supporting a notion that hostile interactions do not always lead to increased aggressiveness between opponents, especially if asymmetric aggression or lack of reciprocal retaliation is in place. Although a common event in nature, conflict can be a limiting factor for species coexisting. While surpassing acceptable thresholds, excessively high levels of aggression can jeopardise relationships between organisms and lead entire colonies to collapse. The behavioural adaptations we described, seem to allow inquilines to manage the amount of aggression received from their hosts. Such a non-threatening individual behaviour may play a fundamental role in cohabitation, as it seems to increase the chances of a stable (although asymmetric) relationship between host and inquiline colonies considerably. We suggest that further research should explore the contributions of such individual actions on collective patterns in the system. While in line with previous reports on cohabitation between termite societies, our findings reinforce the growing view of conflict management as a critical component of socially complex systems. Finally, descriptions of peaceful mechanisms by recipients of aggression in locally restricted, hostile environments should contribute to putting conflict and its consequences in a broader perspective, adding novel insights for studies involving multiple group-living organisms.

## Acknowledgements

We thank Dr Diogro A. Costa, Dr Alessandra Marins and Dr Vinícius B. Rodrigues for useful discussions; Júlio H. Santos and Kátia C. D. Santos for helping with data collection; Dr Fernando Valicente from the Brazilian Enterprise for Agricultural Research (EMBRAPA) and Prof. Ângelo Fonseca for all logistic support. This study was partially funded by (i) the National Council of Technological and Scientific Development (CNPq), (ii) the Foundation for Research of the State of Minas Gerais (FAPEMIG: APQ-08811-15 and BPV-00055-11) and (iii) the DFG Centre of Excellence 2117 “Centre for the Advanced Study of Collective Behaviour” (ID: 422037984). HH was supported with a scholarship from the Foundation for Research of the State of Minas Gerais (FAPEMIG) and a Research grant from the Deutscher Akademischer Austauschdienst (DAAD). PFC holds a Research Fellowship from Coordination for the Improvement of High Education Personnel (CAPES; PNPD no.1680248). ODS holds a Research Fellowship from CNPq (PQ 305736/2013-2). This paper is a contribution from the Lab of Termitology, Federal University of Viçosa, Brazil (http://www.isoptera.ufv.br), deriving from HH’s MSc thesis.

## Author contribuitions

HH, PFC & ODS conceived the study; HH & PFC developed fieldwork, behavioural recording and ethogram design. HH conducted data collection, statistical analysis and drafted the paper. PFC & ODS commented on the manuscript, improving it considerably for submission.

## Ethical statement

We obtained all required permits for the present study, thus complying with relevant regulations governing animal research in Brazil. This includes: (i) permits from The Brazilian Institute for the Environment and Renewable Natural Resources (IBAMA, no. 33094); (ii) permission from The Brazilian Enterprise for Agricultural Research (EMBRAPA) at Sete Lagoas; (iii) permission from landowners at the Divinópolis site to conduct the study on their property; and (iv) tacit approval from the Brazilian Federal Government implied by employing authors to conduct scientific research. None of the sampled species had protected status. No genetic information was accessed in the study.

