## Supplementary material for "Peaceful behaviour: a strategy employed by an obligate nest invader to avoid conflict with its host species"

This is a supplementary material for the paper:

**Contents:**

Table S1; Table S2; Figure S1; Figure S2; Video S1; Video S2; Video S3

**Table S1.** Absolute numbers and percentage of Between-species and Within-species observations annotated from video-samples in closed arenas. Data is presented for hosts (*C. cyphergaster*) and inquilines (*I. microcerus*), and their respective castes.

| Group observed | Number of focal animals (n) | Observations | |
|---|---|---|---|
| | | Between-species | Within-species |
| **HOSTS (*C. cyphergaster*)** | **10** | **199 (32%)** | **421 (68%)** |
| Host workers | 5 | 106 (34%) | 204 (66%) |
| Host soldiers | 5 | 93 (30%) | 217 (70%) |
| **INQUILINES (*I. microcerus*)** | **10** | **125 (20%)** | **495 (80%)** |
| Inquiline workers | 5 | 25 (8%) | 285 (92%) |
| Inquiline soldiers | 5 | 100 (32%) | 210 (68%) |
| **TOTAL** | **20** | **324 (26%)** | **916 (74%)** |

**Table S2** Adjacency matrices containing the behavioural change for each caste of hosts (*C. cyphergaster*) and inquiline (*I. microcerus*). The data was used to extract centrality measures and draw the networks in yED (Abbreviations: walk=walking; rest=resting; aW=antennating wall; aCS=antennating nestmate; aHS=antennating non-nestmate; ig=ignoring; pa=bypassing; rever=reversing; att=attacking).

| Adjacency matrix - Constrictotermes cyphergaster (worker) | | | | | | | | | |
|---|---|---|---|---|---|---|---|---|---|
| | aCS | aHS | aW | att | ig | pa | rest | rever | walk |
| aCS | 4 | 1 | 3 | 0 | 0 | 0 | 0 | 0 | 8 |
| aHS | 1 | 0 | 0 | 3 | 1 | 0 | 0 | 0 | 3 |
| aW | 3 | 1 | 16 | 2 | 1 | 0 | 0 | 0 | 3 |
| att | 3 | 1 | 3 | 11 | 1 | 1 | 0 | 0 | 6 |
| ig | 1 | 1 | 1 | 1 | 0 | 0 | 0 | 0 | 2 |
| pa | 3 | 0 | 1 | 0 | 0 | 0 | 0 | 0 | 2 |
| rest | 0 | 0 | 0 | 1 | 0 | 1 | 1 | 0 | 2 |
| rever | 0 | 1 | 0 | 1 | 0 | 0 | 0 | 0 | 1 |
| walk | 1 | 3 | 3 | 9 | 3 | 3 | 3 | 3 | 26 |

| Adjacency matrix - Constrictotermes cyphergaster (soldier) | | | | | | | | | |
|---|---|---|---|---|---|---|---|---|---|
| | aCS | aHS | aW | att | ig | pa | rest | rever | walk |
| aCS | 2 | 1 | 4 | 1 | 0 | 1 | 0 | 1 | 7 |
| aHS | 0 | 2 | 1 | 0 | 0 | 1 | 2 | 0 | 3 |
| aW | 2 | 0 | 6 | 2 | 3 | 0 | 0 | 1 | 1 |
| att | 2 | 1 | 1 | 1 | 0 | 1 | 1 | 0 | 2 |
| ig | 0 | 2 | 1 | 0 | 2 | 1 | 0 | 1 | 4 |
| pa | 1 | 0 | 0 | 0 | 0 | 0 | 0 | 2 | 3 |
| rest | 2 | 1 | 0 | 1 | 2 | 0 | 15 | 0 | 0 |
| rever | 1 | 0 | 0 | 0 | 0 | 0 | 0 | 1 | 7 |
| walk | 6 | 4 | 3 | 4 | 4 | 2 | 3 | 3 | 24 |

| Adjacency matrix - Inquilinitermes microcerus (worker) | | | | | | | | | |
|---|---|---|---|---|---|---|---|---|---|
| | aCS | aHS | aW | ig | pa | rest | rever | walk | att |
| aCS | 11 | 1 | 3 | 0 | 0 | 9 | 0 | 6 | 0 |
| aHS | 1 | 0 | 0 | 0 | 0 | 1 | 0 | 1 | 0 |
| aW | 1 | 1 | 2 | 1 | 0 | 0 | 0 | 2 | 0 |
| ig | 1 | 0 | 0 | 2 | 0 | 0 | 1 | 0 | 0 |
| pa | 1 | 0 | 0 | 0 | 0 | 0 | 0 | 2 | 0 |
| rest | 7 | 1 | 0 | 2 | 1 | 24 | 0 | 11 | 0 |
| rever | 0 | 0 | 1 | 0 | 0 | 3 | 0 | 1 | 0 |
| walk | 8 | 0 | 1 | 1 | 2 | 12 | 3 | 25 | 0 |
| att | 0 | 0 | 0 | 0 | 0 | 0 | 0 | 0 | 0 |

| Adjacency matrix - Inquilinitermes microcerus (soldier) | | | | | | | | | |
|---|---|---|---|---|---|---|---|---|---|
| | aCS | aHS | aW | att | ig | pa | rest | rever | walk |
| aCS | 5 | 2 | 0 | 0 | 1 | 0 | 0 | 1 | 10 |
| aHS | 0 | 1 | 0 | 2 | 0 | 1 | 0 | 0 | 3 |
| aW | 1 | 0 | 0 | 0 | 0 | 1 | 0 | 0 | 2 |
| att | 0 | 1 | 0 | 12 | 0 | 0 | 0 | 2 | 5 |
| ig | 0 | 0 | 1 | 0 | 1 | 0 | 0 | 1 | 4 |
| pa | 0 | 1 | 0 | 0 | 0 | 0 | 2 | 2 | 4 |
| rest | 0 | 0 | 0 | 0 | 1 | 0 | 4 | 0 | 3 |
| rever | 2 | 0 | 2 | 0 | 0 | 3 | 0 | 2 | 5 |
| walk | 11 | 2 | 1 | 6 | 4 | 4 | 1 | 6 | 27 |

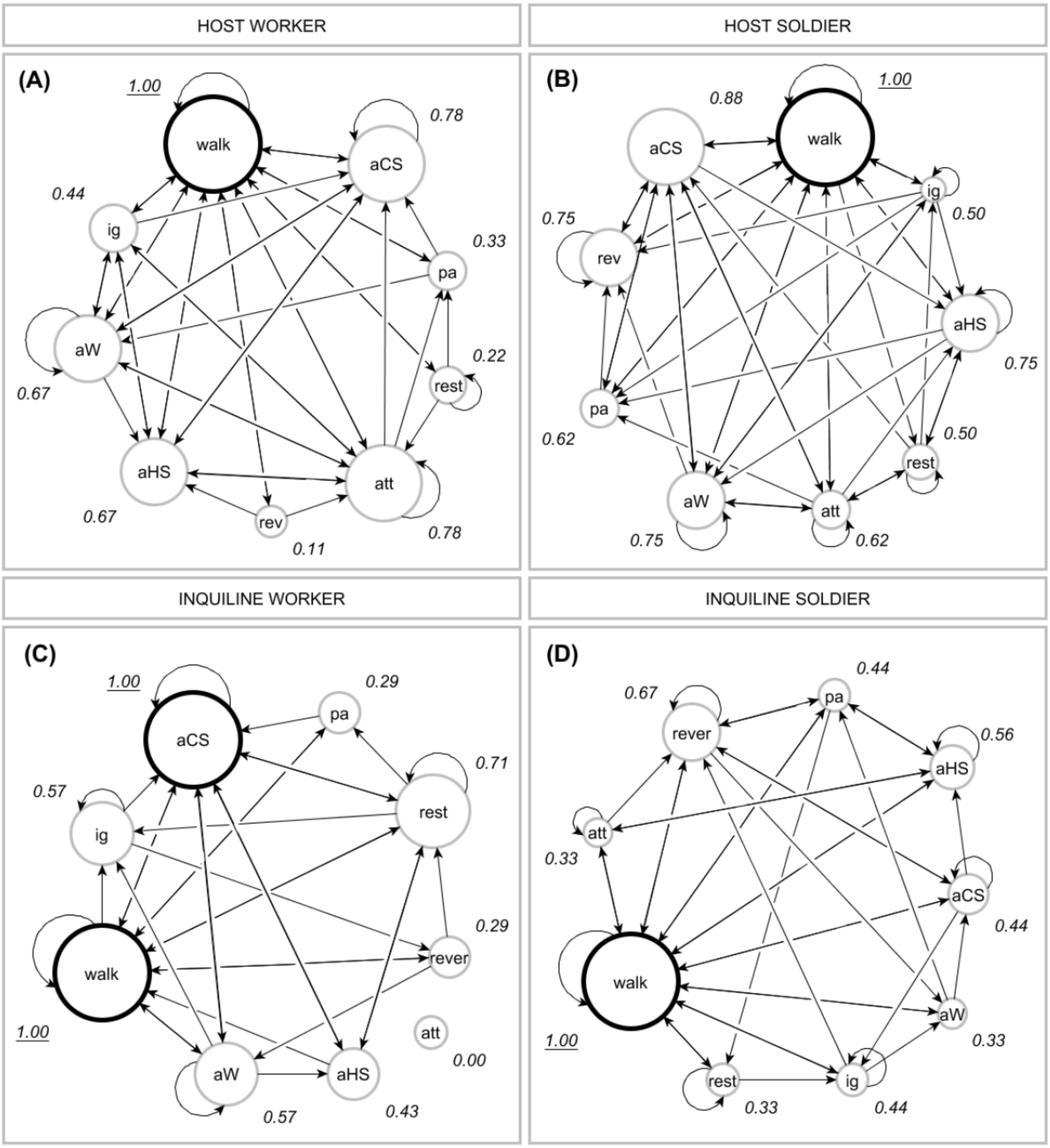


**Figure S1.** Behavioural profiles observed for each caste (with scores of centrality measure). Nodes represent behaviours performed by individuals, whereas connecting edges (arrows) represent behavioural changes occurred from one behaviour to another. Behaviours with the highest influence othe n network are highlighted with thicker node contours. Node size was adjusted using calculated centrality measures (scores) to visually represent the degree of influence exerted by each behaviour upon the profiles. (Abbreviations: walk=walking; rest=resting; aW=antennating wall; aCS=antennating nestmate; aHS=antennating non-nestmate; ig=ignoring; pa=bypassing; rever=reversing; att=attacking).

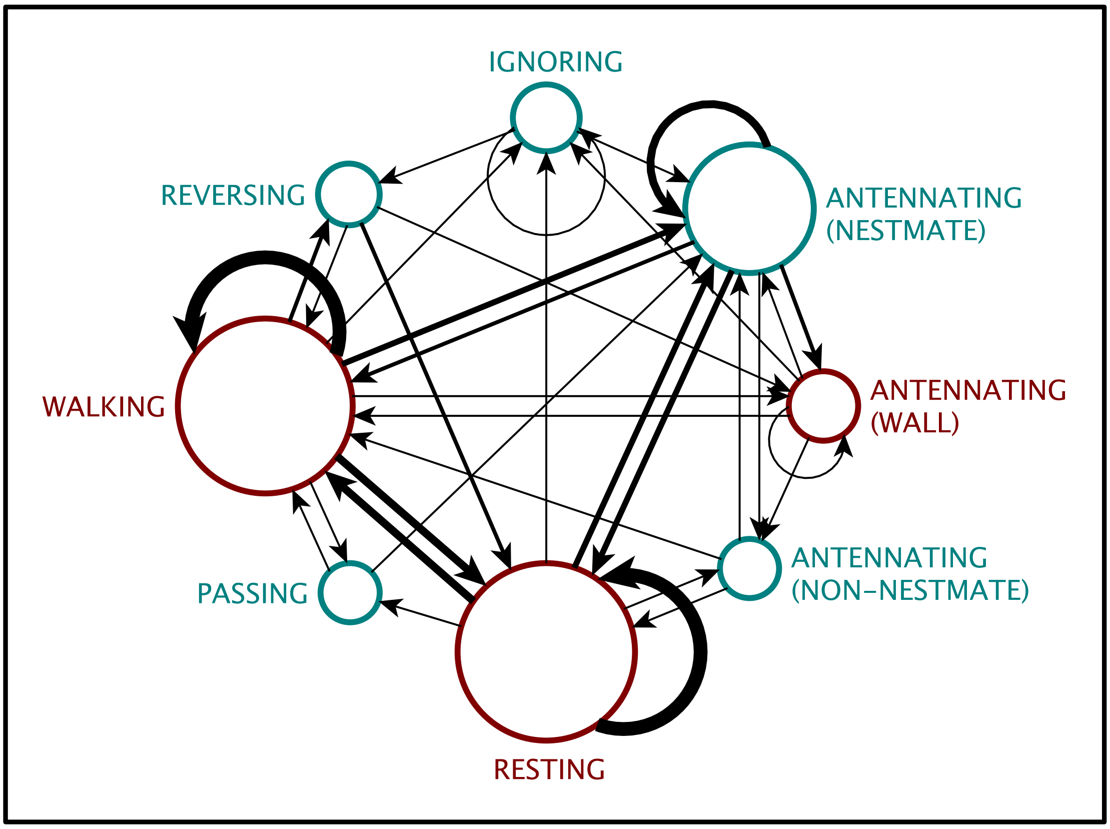


**Figure S2.** Behavioural change for inquiline workers: nodes represent behaviours performed by individuals, whereas connecting edges (arrows) represent behavioural changes occurred from one behaviour to another. Node size was adjusted using behavioural frequencies extracted from the annotation. The width of edges indicate how frequently a given behavioural change occurred.

**Video S1**

**S1_snapping.mov**

This video was recorded in lab conditions with a fixed individual of *Termes* sp.. It shows the mechanism of snapping, also present in *I. microcerus* and other termite species with soldiers provided of slender mandibles.

**Video S2**

**S2_snapping2.mov**

This vídeo was recorded in lab conditions with a free individual of *I. microceus* in the presence of *C. cyphergaster*. It shows snapping events performed by the inquiline soldier of in retaliation to host threats.

**Video S3**

**S3_defecation.mov**

This vídeo was recorded in lab conditions with individuals of *I. microceus* and *C. cyphergaster* in experimental arenas. It shows the aggressive nature of host-inquiline encounters and some of the evasive behaviours performed by inquilines as response. Markers were included to highlight when aggressive interactions happened. The defensive mechanism using defecation is the depicted in the footage with several events.
